# PLGA-based dual-loaded nanoformulation of DIM and TMZ - An advanced clinical strategy for brain cancer treatment in a combinatorial approach

**DOI:** 10.1101/2023.05.04.539373

**Authors:** Sibani Sarkar, Sunny Kumar, Gouranga Saha, Malini Basu, Mrinal K. Ghosh

## Abstract

Glioblastoma multiforme (GBM) is a highly aggressive form of primary brain tumor in adults, which unfortunately has an abysmal prognosis and poor survival rates. Even though several FDA-approved multimodal treatments for targeting GBM are available, the effectiveness in most patients are not satisfactory. The reason behind this poor success rate is mainly attributed to insufficient drug distribution to the tumor site across the blood-brain barrier (BBB) and induction of resistance for single-drug based therapies. Chemotherapy with Temozolomide (TMZ) having a median overall survival of around 12-15 months, envisages the urgent necessity for more effective treatment strategies. Based upon these facts, in this study, we have developed a novel approach for repurposing TMZ along with inhibition of EGFR, which overexpressed in GBM, to achieve our goal. PLGA-based nanoencapsulation of both TMZ and 3,3’-diindoyl methane (DIM), an EGFR inhibitor, in a combinatorial approach enhances the delivery of them together. Their synergistic mode of actions, significantly enhances the cytotoxic effect of TMZ *in vitro* and *in vivo*. Moreover, the dual-loaded nanoformulation works more efficiently than their individually packed nanoparticles on DNA damage and apoptosis, resulting in a several-fold reduction in tumor burden, systemic drug toxicity, and increased survival. These findings suggest the preclinical potential of this new treatment strategy.

## Introduction

Glioblastoma, or grade IV astrocytoma, is one of the deadliest and most aggressive types of adult brain tumors. Clinical data shows that GBM has a poor prognosis showing a survival time of 12–15 months, and less than 5% of patients survive five years after diagnosis. GBMs can arise in the brain either de novo or evolve from lower-grade astrocytoma. In adults, GBM occurs most often in the cerebral hemispheres, especially in the frontal and temporal lobes of the brain ^1^. A glioma patient underwent standard clinical strategies, including maximum surgical removal of the tumor followed by chemo-and radiotherapy, even though the overall survival of GBM patients has barely prolonged. Several factors like BBB, development of chemoresistance, glial stem cell population, acquired radioresistance, and tumor heterogenicity significantly reduced the effectivity of the chemotherapeutic drugs used and increased recurrence ^2^.

TMZ has been widely used as a standard chemotherapeutic option in treating newly diagnosed GBM since its initial FDA approval in 2005. In the landmark paper, Stupp *et al*., demonstrated that the addition of TMZ to radiotherapy resulted in an increased median survival of up to two months, which remains one of the most significant improvements in GBM treatment to date ^3^. But further research established some major concern associated with TMZ-based glioma treatment is the development of TMZ resistance. Unfortunately, over 50% of GBM patients show resistance to TMZ due to the highly heterogeneous and mutation-prone nature of GBM. It induces cytotoxicity by DNA methylation of guanine and adenine residues ^4^. Specifically, the alkylation of the O6 position of guanine leads to nucleotide mispairing during DNA replication, resulting in single and double-strand DNA breaks, cell cycle block, and ultimately, apoptosis ^5–7^. GBM patients can develop resistance to TMZ due to several factors, including higher expression of MGMT that reverses methylation of guanine, epigenetic regulation, and autophagy induction. Higher concentrations of TMZ may be necessary to overcome resistance, but this can lead to toxic side effects ^8–12^. Furthermore, resistance to single therapies can develop resistance due to the genetic heterogeneity and adaptability of GBM. Identifying rational and effective combination therapies has become increasingly important in addressing this challenge. Combining anticancer drugs can enhance efficacy compared to monotherapy by targeting pathways in a cooperative or additive manner, potentially reducing drug resistance and providing therapeutic anticancer benefits, such as reducing tumor growth and metastatic potential. However, few established combination therapy trials exist, and their potency has not yet been thoroughly investigated ^13^.

DIM is a phytochemical that can be a potential chemotherapeutic agent and is produced from the digestion of Indole 2 carbinol (I3C) in cruciferous vegetables ^14, 15^. DIM has shown anti-tumor effects in preclinical models, and it targets multiple aspects of cancer progression, such as cell cycle regulation and survival, through modulating EGFR-JAK2-STAT3 signalling, checkpoint activation, caspase activation, ER stress generation, autophagy, and anoikis. DIM has low systemic toxicity, and it reduces carcinogen-induced breast and lung tumor formation with reduced toxicity ^16^. DIM also has the ability to sensitize the cancer cell to standard chemotherapeutic agents and can reduce the toxicity and drug resistance associated with available chemotherapeutics like TMZ. DIM is widely used in combination therapy to generate chemosensitization and reduce the toxicity of the accompanied drug ^17^.

Nanotechnology has emerged as a promising platform for cancer therapy by enabling precise targeting and delivery of therapeutics to the tumor site. Nanoparticles offer a safe and efficient means of delivering drugs to the tumor region. Significant clinical responses can be achieved using a BBB-penetrating polymeric nanoparticles that are taken up by GBM cells and release the encapsulated drug(s) over time. Conclusively, nanoparticles represent a highly effective approach for cancer treatment, providing unparalleled precision and efficacy. Among the variety of materials used for the fabrication of nanosystems, Poly (lactic-co-glycolic acid) (PLGA) has gathered special attention due to its biocompatibility and biodegradability, and widely used for effective tumor targeting by efficient drug delivery to the tumor site. PLGA-based nanoparticles can be used to deliver small chemicals, peptides, proteins, and nucleic acids used in cancer therapy ^15, 18^. PLGA nanoparticles improved drug permeability and retention effect (EPR) by attaching nano molecules to the tumor cells surface for efficient diffusion ^19^.

In this study, we chose PLGA for the nanoencapsulation of DIM and TMZ over other materials because it improves sustained release, biocompatibility, biodegradability, and efficiency of delivery across the BBB. Additionally, the nanoformulations in a combinatorial approach or dual-loaded formulation induced glioma cell death by activating the caspase dependent pathway and showed anti-tumor effects by inducing apoptosis of the glioma cells in the tumor-bearing rat. Briefly, our results emphasize the effective delivery of DIM and TMZ to combat glioma progression by activating programmed cell death pathways.

## Methods

### Cell culture and reagents

Rat glioma C6 cells were obtained from ATCC (American Type Culture Collection) and National Centre for Cell Science (NCCS). Cells were cultured in DMEM supplemented with 10% Fetal Bovine Serum (FBS) and 1% penicillin/streptomycin in 5% CO2 at 37℃. C6 cells were treated with nanoencapsulated DIM and TMZ either alone or in combination or as dual-loaded nanoformulation for indicated time points ^20^.

### Cytotoxicity study

MTT assay was performed on C6 cells in ninety-six well dishes after treatment with native DIM, TMZ or with their PLGA nanoformulations for 24 hrs keeping DMSO or empty nanoformulation as control. The assay was performed using an established protocol described previously and readings were taken in triplicates at 570 nm using Ultra Multifunctional Microplate Reader ^15^.

### Synergy Calculation

Compusyn software (ComboSyn, Inc.) was used to assess the synergistic, additive, and antagonistic effects of DIM and TMZ, either native states or as nanoformulations, using the Chou-Talalay method for drug combination. The combination index (CI) value was calculated as CI=(D1/DX1) + (D2/DX2), where D1 and D2 denote the doses of agents 1 and 2, respectively, while DX1 and DX2 denote the doses of single agents 1 and 2 under a certain proliferation inhibitory rate. CI<1 indicates a synergistic effect, while CI>1 indicates an antagonistic effect ^21–23^.

### Bioinformatics based ADME prediction

Several ADME properties such as physicochemical (Mol. weight, TPSA), pharmacokinetic (skin (LogKp), gastrointestinal (GI) and BBB permeability, solubility (ESOL, Ali), drug likeliness (lipinski, bioavailability score), lipophilicity (iLogP, wLogP) of native DIM and TMZ were predicted using the SwissADME based online server (http://www.swissadme.ch/)^24^.

### Bioavailability radar model based prediction

By using canonical SMILES of native DIM and TMZ, the oral bioavailability of these drugs were predicted from six physicochemical properties such as lipophilicity, size, polarity, insolubility, in-saturation and flexibility of radar model ^24^.

### Boiled-egg model based brain vs. GI penetration prediction

Boiled-Egg model obtained from SwissADME online server, a plot between wLogP *vs* Topological polar surface area (TPSA) were used to determine the penetration of native drugs *viz.,* DIM and TMZ through BBB and GI ^24^.

### Development of PLGA-based nanoformulation

PLGA-encapsulated nanoparticles were prepared using a modified emulsion diffusion evaporation method. 1mg each DIM and TMZ for individual or 1:2.5 ratio (DIM:TMZ) for dual nanoformulation were dissolved in 1 ml ethyl acetate, and 50 mg PLGA was dissolved in 5 ml ethyl acetate. The two solutions were mixed and emulsified by dropwise addition to water containing DMAB, then stirred for 3 hrs at room temperature until the solvent evaporated. The resulting nanoparticles were washed, resuspended in PBS, and lyophilized before being freshly used or stored at –20°C ^15, 25^.

### Characterization of the nanoformulations

*Dynamic Light Scattering (DLS):* The particle size of all the prepared nanoformulations were measured by Nanosizer 90 ZS (Malvern Instruments, Southborough, MA). 1 mg of lyophilized powder was suspended in 10 ml water and passed through a 0.22 µM filter. The size was measured as represented by an average value of 20 runs ^15, 25^.

### Transmission Electron Microscopy (TEM)

The size and shape of the nanoformulated compounds were determined in dry state using a JEOL-JEM 2100 TEM (JEOL, Inc., Peabody, MA, USA, acceleration voltage of 300KV). The lyophilized powder was suspended in Milli Q water and mounted on a copper grid (300 meshes) with a thin film of amorphous carbon ^26^.

### Atomic Force Microscopy (AFM)

Nanoparticle’s size and shape was measured by 5500 Pico Plus AFM system, Agilent Technologies. The lyophilized powder was suspended in Milli Q water and poured onto a mica sheet. After drying, the nanoparticle’s size was determined using cantilevers with specific resonance frequency (146–236 kHz), tip height and length (10–15 μm, and 225 μm, respectively), and imaging was done in the aquatic mode ^15, 27^.

### Zeta Potential

To understand the surface charge of nanoparticles and their stability, the Zeta potentials of the synthesized nanoparticles were measured by Zetasizer Nano ZS (Malvern Instruments, Southborough, MA) ^15^.

### Entrapment efficiency and in vitro release study

A proportion of lyophilized particles was suspended in ethyl acetate and shaken for 3 hrs. The entrapment efficiency was measured spectrophotometrically using UV-VIS spectrophotometer model Schimatzu (Singapore). The drugs were detected at their specific absorbance frequencies. The EE parameter was calculated as EE = (Wt-Wd)/Wt (where Wt and Wd describe the total drug added and extracted into the supernatant, respectively). For release assay, the nanoparticles were incubated in PBS, followed by centrifugation. The amount of released drug in the supernatant was periodically determined using a spectrophotometer for up to 72 hrs ^27^.

### FTIR study

Encapsulation of the compounds was further studied by Fourier Transform Infrared Spectroscopy (FTIR) using a model spectrum TWO (Perkin Elmer, USA). The FTIR spectra of native compounds and their nanoformulations were obtained by mixing samples with KBR powder (Infrared grade) ranging from 400-4000 cm^-1^ ^15, 25^.

### Measurement of Mitochondrial Membrane Potential (MMP) and Reactive Oxygen Species (ROS)

C6 cells were exposed to native or nanoformulated compounds for 24 hrs followed by staining with JC-10 dye for 1 hr. The fluorescence of JC-10 aggregates and monomers were measured at an excitation/emission wavelengths of 490/590 nm ^28^.

The intracellular ROS levels in C6 cells were determined using the Fluorometric Intracellular ROS Kit (MAK142; Sigma, USA), which was performed in a 96-well plate format with detection at an excitation/emission wavelengths of 650/675 nm ^29^.

### Flow Cytometry

C6 cells were treated with indicated nanoformulations for fixed time points followed by Annexin V/ propidium iodide (PI) staining using Annexin V-FITC Apoptosis Detection Kit II (Cat # PF032, Sigma-Aldrich), as per manufacturer protocol. Data were obtained using LSR Fortessa (BD Biosciences) and analyzed using FlowJo Software ^30, 31^.

### Immunoblotting

C6 cells were treated with specified nanoformulations or native drugs for 24 hrs before being harvested in lysis buffer. 30 µg of proteins were then subjected to immunoblotting using primary and HRP-tagged secondary antibodies. The membrane was washed with TBST and bands were detected using the ECL detection system ^32, 33^.

### Cytochrome C Release Assay

C6 cells were seeded on poly-L-lysine coated coverslips and treated with the indicated nanoformulations. The treated cells were immunostained with Cytochrome C and HSP60 antibodies, and colocalization was studied using standard immunocytochemistry protocol. Cells were observed under a confocal microscope, and captured images were processed using Image J software ^34^.

### Soft agar colony formation and two-dimensional scratch assays

Cells were mixed in 0.35% complete agarose medium and plated on the bottom layer in 35mm plates containing 0.7% agarose medium. The culture media containing different nanoparticles were changed every alternative day for 14 days. Colonies were counted in three microscopic fields using Olympus IX-81 microscope and captured using image pro plus imaging software ^31^.

For the scratch assay, 5000 cells were seeded on 35 mm plates and grown to reach required confluency before indicated treatments. Scratches were made using 10 µl sterile pipette tips. 24 hrs post-treatment, the migration of cells was observed microscopically at premarked positions ^35^.

### Animal Maintenance

Adult SD rats (120 gms, 30-45 days old) were obtained from the animal house of the Indian Institute of Chemical Biology (IICB) and maintained under optimum conditions (12:12 dark:light circles) with standard pellet diet and water ad libitum. All experiments were conducted following CPCSEA guidelines and approved by the animal ethics committee of CSIR-Indian Institute of Chemical Biology (IICB) ^20, 32, 33^.

### Intracranial brain tumor model

Sprague-Dawley rats (150-160 gms) were anesthetized and intracranially injected with early passaged C6 cells (10^6^) using stereotaxis according to previously standardized protocol. Tumor formation was detected after 14 days by observing behavioral and physiological features such as hair loss, movement difficulties, aggression and weakness. Treatments were initiated immediately in randomized groups after tumor formation. After treatment for the indicated time points, the tumor-bearing rats were sacrificed by cervical dislocation, and the brains were collected to examine the tumor status. A portion of each tumor was fixed in 10% buffered formalin, embedded in paraffin for block preparation followed by histology and IHC analysis of tissue sections ^15, 32, 36^.

### Subcutaneous tumor model in flank region of rats

C6 cells were injected in the flank region of female SD rats (19). After 12 days, the tumor developed in the flank region was detected by palpating and noticing behvioural changes. Treatments were started immediately after tumor development and continued for up to 40 days while every alternate day, tail-vain injection of nanoformulations of indicated doses was done ^31–33^.

### Histology and Immunohistochemistry (IHC)

Tumor tissues were fixed in 10% neutral buffered formalin, embedded in paraffin, and cut into 5 µm sections using a Microtome. Hematoxylin and eosin staining was used to visualize tissue morphology, while IHC was performed using a citrate buffer for antigen retrieval, blocking with serum, incubation with primary antibodies overnight at 4°C, and HRP-tagged secondary antibody for 2 hrs at room temperature. The final step involved observing DAB-stained sections under the microscope and captured the images ^36, 37^.

### TUNNEL Assay

After treating with indicated nanoformulations for 24 hrs, cells were fixed with 4% formaldehyde for 15 min on ice, followed by a PBS wash. The resulting cell pellet was incubated in 70% ice-cold ethanol for 30 min, washed twice in a wash buffer, and then resuspended in staining solution for 60 min at 37°C as per the manufacturer’s protocol (TUNNEL Assay Kit – FITC; cat # ab66108). After rinsing, the TUNNEL-positive cells were detected by FACS. A modified protocol was used to detect TUNNEL-positive cells in tumor tissue samples. The sections were deparaffinized, rehydrated, and stained with a staining solution for 60 minutes at 37°C. The sections were washed twice, mounted on DAPI-containing mounting media, and visualized under a fluorescence microscope ^32^.

### Statistical analysis

All statistical calculations were done using GraphPad Prism 8.4.2, and H scores of IHC intensities performed using Image J software. Statistical analysis was performed by Student’s t-test, or one-way ANOVA, as reported previously.

## Results

### Native DIM and TMZ act synergistically to induce apoptosis in glioma cells

To evaluate the chemotherapeutic potential of the drugs chosen for this study, we conducted several *in vitro* assays on GBM cells. First, we examined whether DIM could enhance the anti-tumor activity of TMZ when used together. We treated C6 glioma cells with either DIM or TMZ for 24 hrs and then determined the IC50 values using the MTT assay. The results revealed that both DIM and TMZ had a dose-dependent ability to inhibit the growth and induce apoptosis of C6 cells. Their respective IC50 values were 32.3 ± 2.54 µ M and 601 ± 3.61 µM (**Fig. 1A**). We then evaluated the combinatorial effect of both the drugs on C6 cells by treating them with multiple dose combinations. We use Chou-Talalay’s method to find the synergistic drug concentrations, if any. Surprisingly, our results showed that the Combination Index (CI) value was less than 1 at several drug concentrations chosen, but not in all combinations. We identified that the CI value was the lowest at 12.5 µM DIM and 500 µM TMZ among all the CI values that we determined and selected this combination for subsequent experiments (**Fig. 1B**). To determine the effect of combining DIM and TMZ at their synergistic concentrations on cell death induction, we further monitored reactive oxygen species (ROS) production following treatment with the drugs alone or in combination. Our results show that the combined treatment produced significantly more ROS than the individual compounds or vehicle control (**Fig. 1C**). The intensity of JC10 orange J-aggregate emission profile decreased by approximately 50% in C6 cells treated with either DIM (12.5 µM) or TMZ (500 µM) used alone, and by approximately 65% in cells treated with the synergistic combination of both compounds, indicating a significant loss of mitochondrial membrane potential (MMP) (**Fig. 1D**). This decrease in MMP, coupled with increased ROS production, resulted in increased stress and a reduction in cell number after 24 hrs of treatment, as confirmed by bright-field microscopy (**Fig. 1E**).

**Fig. 1.**
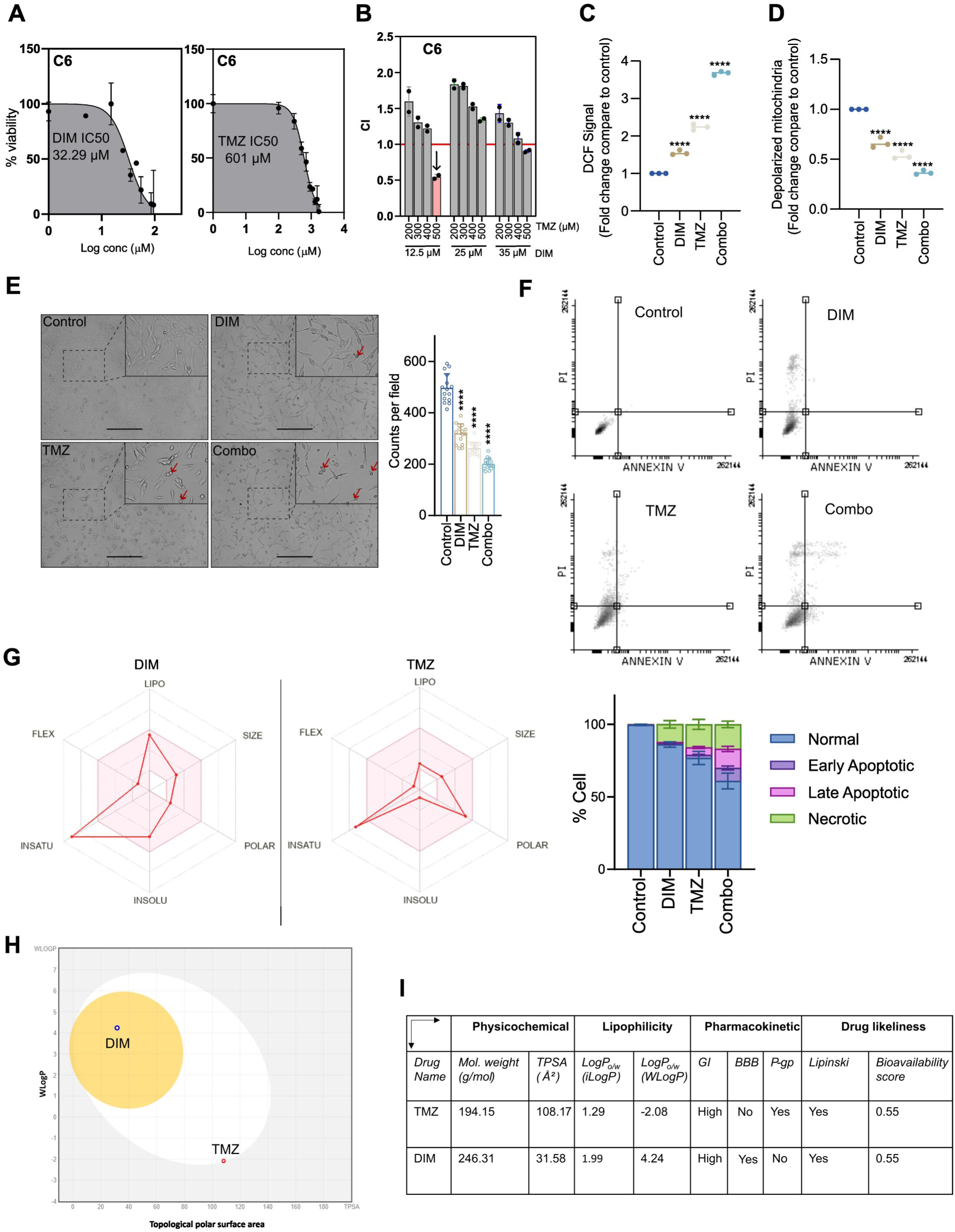
Toxicity study of native DIM and TMZ on Glioma cells. **A.** *In vitro* cytotoxicity study of DIM and TMZ. **B.** Synergestic mode of actions of DIM and TMZ by MTT assay. **C.** ROS generation in control, DIM, TMZ and their combination use using a fluorescein-labelled probe DCFH. **D.** Flow cytometric evaluation of MMP using JC-1 as molecular probe. **E.** Changes in morphology and density of cultured C6 cells after treatment with DIM and TMZ either alone or in combination using Bright Field Microscopy. **F.** Flow cytometric detection of percent (%) apoptosis of control, nDIM, nTMZ and combination of DIM and TMZ treated C6 cells using Annexin V-FITC staining. **G.** Bioavailability radar model, **H.** Boiled-egg model and **I.** SwissADME properties of native DIM and TMZ as depicted in the figure.

We further estimated the apoptotic cell populations of C6 cells by Annexin V/PI-FACS analysis to emphasize the cytotoxic effect of the synergistic dual treatment. Compared to the DMSO control, DIM and TMZ treatment increased the number of apoptotic cells by ∼14% and ∼18%, respectively, and their combined treatment led to a significant increase in the number of apoptotic cells by ∼39% after 24 hrs. In C6 glioma cells, the DIM and TMZ combination treatment efficiently induced a noticeable increase in early and late apoptotic cell populations compared to control or their individual treatments, suggesting its superior efficacy in inducing apoptosis when used them together. Treatment with the individual drugs resulted in severe cytotoxicity, characterized by a higher population of necrotic cells without any significant percent of early or late apoptotic cells, which was reduced by combination therapy, underscoring the efficacy of the combined treatment in inducing cell death with significant increase in early and late phase population of apoptotic cells (**Fig. 1F**). Thus, further studies are required to critically investigate the underlying mechanisms of this synergistic effect, and combination therapy may be more effective for treating C6 rat glioma cells than their individual treatments.

Several bioinformatics-based pharmacokinetics studies were conducted to determine the bioavailability of these two drugs. The radar model revealed that DIM and TMZ are not orally bioavailable in their native form due to their high saturation, resulting in poor solubility in the human gastrointestinal tract and low bioavailability (**Fig. 1G**). Additionally, the Boiled-egg model predicted that DIM is highly penetrable and TMZ is partially penetrable to the brain (**Fig. 1H**). Furthermore, SwissADME-based properties indicate that TMZ is effluxed by P-gp, leading to a lower concentration in the brain region (**Fig. 1I**). All these findings suggest nano-encapsulation of DIM and TMZ may improve their synergistic actions in glioma therapy by minimizing the required dose.

### Characterization of individually packed DIM and TMZ as PLGA nanoformulations

To achieve our goal, we further planned to prepare individually packed nanformulations of DIM (nDIM) and TMZ (nTMZ) using FDA-approved PLGA for their encapsulation and used them either individually or in combination. We aimed to address two common issues with glioma therapy: i) to reduce the cytotoxicity of drugs and ii) to improve the passage of these drug molecules across the BBB. We used a modified emulsion-diffusion evaporation method to encapsulate DIM and TMZ in PLGA. In our experimental setup, the drug-polymer ratio of 1:5 produced the highest entrapment efficiency of the nanopreparations (nDIM 85±1.2% and nTMZ 90±2.1%). Furthermore, DMAB enhances the effectivity and successful entrapment of DIM and TMZ in nanovesicles. Additionally, the PLGA-nanoparticles released drug molecules in a controlled and effective manner, indicating robust encapsulation. We observed 10% TMZ from nTMZ and 15% DIM from nDIM were released from their nanoparticles till 80 hrs, maintained sustained release (**Fig. 2A**). The zeta potential values of unloaded (Empty), nDIM and nTMZ nanoparticles were examined and found 12.6±5.3mV, –52.6±14.7mV, and –41±10.6mV, respectively, suggesting enough stability of these nanoformulations (**Fig. 2B**). Additionally, DLS, AFM and TEM analysis of an aqueous dispersion of nDIM, nTMZ and Empty nanoparticles show that all of the particles had a size distribution range of 20–50 nm and were more or less spherical in shape (**Fig. 2C, D & E**). According to AFM analysis, Empty, nDIM, and nTMZ had respective mean widths of 21.24±4.9, 20.44±4.9, and 24.06±5.4 nm, respectively (**Fig. 2D, bar graph**). These characterizations of the prepared nanoformulations suggest that they are stable, easily soluble in water, and can be stored at room temperature and 4°C without experiencing any decay or aggregation for an extended period.

**Fig. 2.**
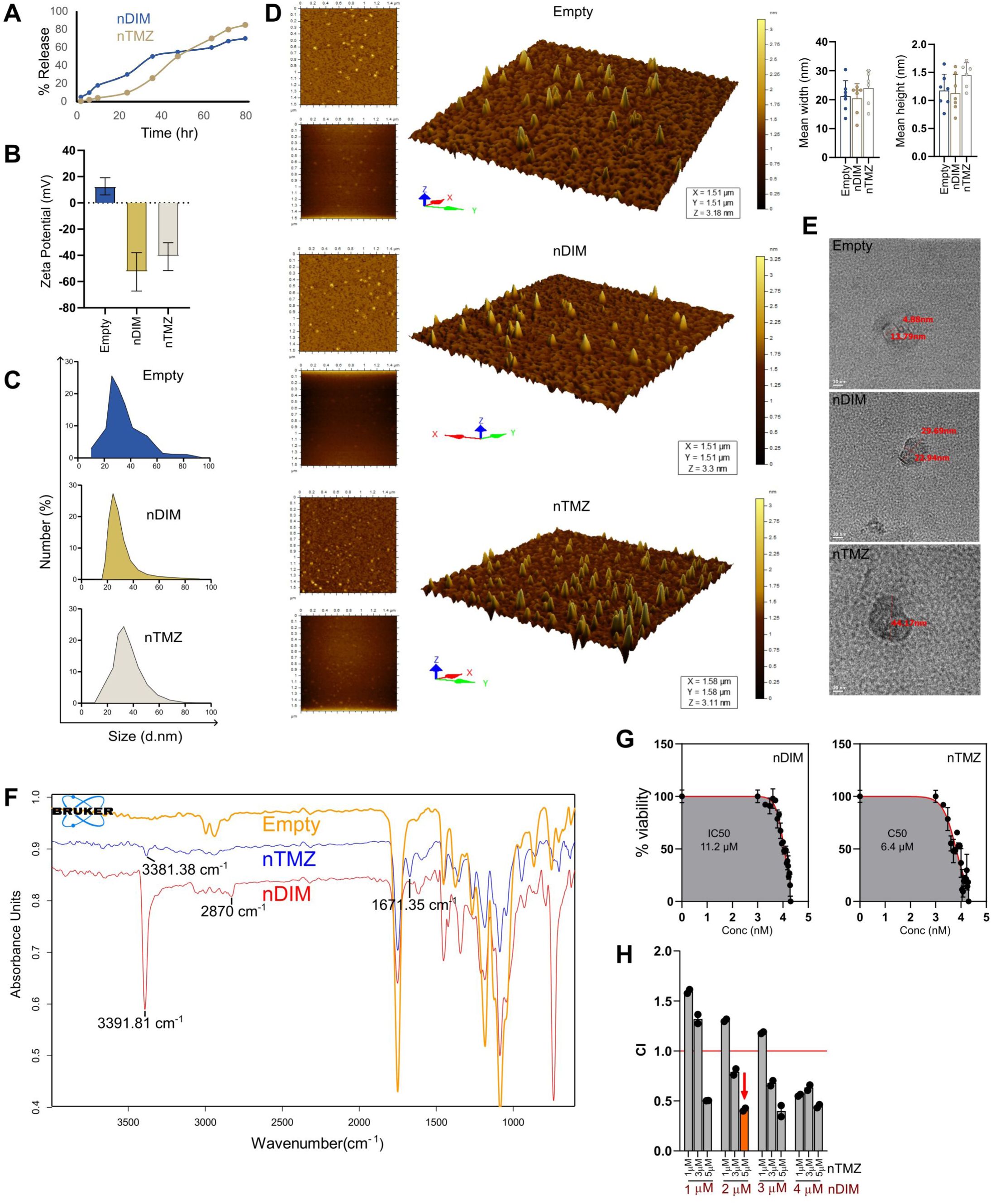
Characterization of individually loaded nanoparticles. **A.** Kinetic release from its corresponding PLGA nanoformulations. **B.** Zeta potential, **C.** Size distribution by DLS of empty, nDIM and nTMZ nanoparticles. **D.** AFM study, **E.** TEM images, **F.** FTIR spectra and **G.** cytotoxicity determination using MTT assay of empty, nDIM and nTMZ nanoparticles. (H) Synergy calculation of nDIM and nTMZ using MTT assay.

Next, we verified the qualitative presence of DIM and TMZ in the nanoformulations using Fourier Transform Infrared (FTIR) spectroscopy. We observed characteristic and distinguished peaks in both nDIM (3391.81 cm^-1^ for amine stretching and 2870 cm^-1^ for C-H stretching) and nTMZ (3381.38 cm^-1^ for amine stretching and 1671.35 cm^-1^ for carbonyl stretching), confirming their presence in the respective nanoformulations (**Fig. 2F**). Furthermore, we also confirmed the presence of the native drug molecules inside the respective nanovesicles by mass spectrometry. Peaks for DIM (247.1228 *m/z*) and TMZ (M+H = 195.0622 *m/z* and M+Na = 217.0441 *m/z*) were observed, confirming their presence in the respective nanoformulations (**Supplementary Fig. 1**). Next, to examine the effect of these nanoformulations on cell viability, we treated C6 glioma cells with nDIM and nTMZ for 24 hrs and observed significantly lower IC50 values of nDIM and nTMZ are 11.2 µM and 6.4 µM, respectively, compared to DIM and TMZ were 32 µM and 601 µM, respectively, indicating that the nanoformulations are much more effective in terms of cytotoxicity than their respective native forms (**Fig. 2G**). The “Dose-CI plot” shows the synergistic cytotoxic effect of the combination therapy, where most of the chosen dose combinations exhibit synergistic activities in terms of induced cytotoxicity. Furthermore, it can be predicted that nDIM effectively increases the sensitivity of C6 cells towards nTMZ. We selected the effective combination with the lowest drug dose and the highest CI-value, which is 2 µM of nDIM and 5 µM of nTMZ, to conduct our subsequent experiments (**Fig. 2H**). The precise mechanism of increased cytotoxicity by these nanoformulations is not fully explainable but it is believed that it is caused by an increased cellular uptake of the drug molecules encapsulated in the nanoformulations. Additionally, the synergistic combination of nDIM and nTMZ enormously decreased the therapeutic dose of TMZ and suggests a novel way to repurpose against glioma. This suggests that using DIM in combination with TMZ as nanoformulations can enhance the efficacy of chemotherapy and reduce TMZ side effects, making them a promising approach for glioma treatment.

### Preparation and characterization of dual-loaded PLGA nanoformulation of DIM and TMZ

To further resolve the issue, we immediately developed dual-loaded nanoformulation to entrap both DIM and TMZ together with the possibility to further improve the efficacy and reduce toxicity. This approach also minimizes errors in handling the individually packed molecules, which could occur when using a mixture of nDIM and nTMZ. We used 1:2.5 molar ratio of DIM and TMZ to create the dual-loaded PLGA nanoparticles, kept in mind use of their minimum concentrations for achieving the highest CI value. We further observed that he dual-loaded nanoparticles maintained the synergistic drug ratio during *in vitro* release, with both DIM and TMZ being released slowly in a sustained manner over a period of several hrs. For instance, within the first 24 hrs, only approximately 20% of DIM and 10% of TMZ were released, while around 75% of both molecules were gradually released over 100 hrs. This controlled release pattern ensures that the therapeutic effects of the drug molecules are sustained over a longer period, improving their overall effectiveness (**Fig. 3A**).

**Fig. 3.**
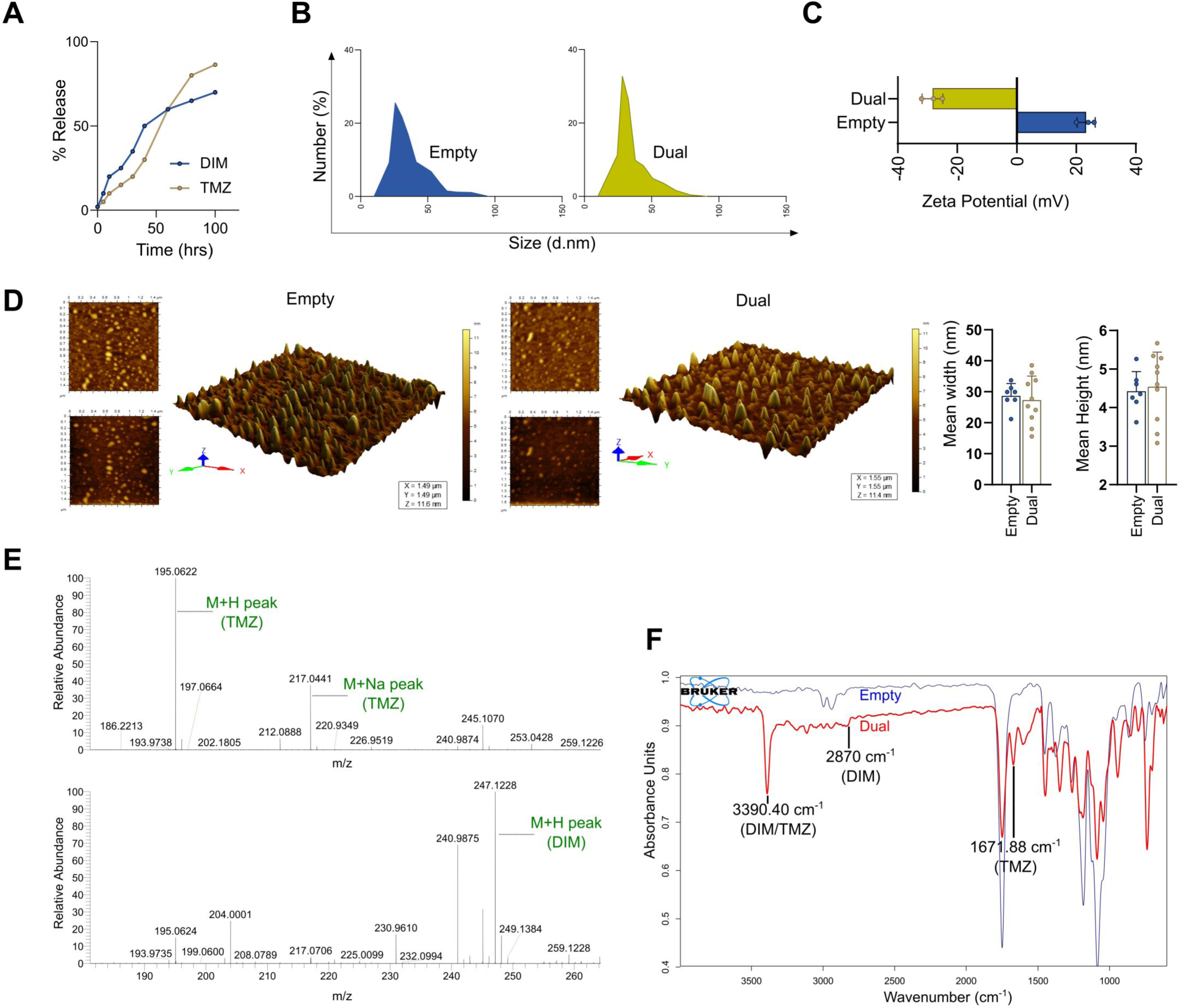
Physical characterization of dual-loaded nanoformulation. **A.** Kinetic release of DIM and TMZ from dual-loaded nanoformulations. **B.** Size distribution using DLS and **C.** Zeta potential of empty and dual-loaded nanoparticles. **D.** Size determination by AFM, **E.** Confirmation of DIM and TMZ nanoformulations. **F.** FTIR spectra of the respective nanoformulations used.

Next, the size of the dual-loaded nanoparticles were determined by DLS and AFM, indicating that their aqueous dispersions were more or less spherical, and the size distribution range was 30±5 nm. The zeta potential values of empty and dual-loaded nanoparticles were 23±2.8 and –25mV±5.8, respectively, indicating good stability (**Fig. 3B, C, and D**).

Moreover, the mass spectrometry plot confirmed the presence of both DIM and TMZ in the dual-loaded nanoparticles as detected by peaks specific to their native forms. The M+H base peak (195.0622 m/z) and M+Na adduct peak (217.0441 m/z) were observed, confirming the presence of TMZ. The M+H peak for DIM was also observed at 247.1228 m/z (**Fig. 3E**). Additionally, FTIR was used to further confirm the presence of both DIM and TMZ in the dual-loaded PLGA nanoparticles. A deep and common amine stretching peaks at 3390.40 cm^-1^, a C-H stretching peak at 2870 cm^-1^ (DIM), and a carbonyl stretching peak at 1671.88 cm^-1^ (TMZ) were observed, indicating the presence of both DIM and TMZ in the dual-loaded nanoparticles (**Fig. 3F**).

### Biological evaluation of dual-loaded nanoformulation in vitro

Anticancer drug treatments like TMZ are known to induce apoptotic signaling pathways in cancer cells by causing DNA damage response, partly due to the production of ROS and mitochondrial depolarization. In this study, we measured ROS generation in treated C6 cells using DCFH-DA staining to assess changes in intracellular redox status. The results showed that the dual-loaded nanoformulation was much more effective in generating ROS than the individually packed nanoparticles. The DCF signal was significantly increased when compared to either nDIM (2µM) or nTMZ (5µM) treatments, indicating that the combination of both drugs as dual-loaded PLGA nanoformulation enhanced ROS production in C6 cells (**Fig. 4A**). Similarly, upon treatment with dual-loaded nanoformulation, a significant level of mitochondrial membrane depolarization was observed in C6 cells, where individually packed molecules were kept for comparative analysis (**Fig. 4B**). The results show that at the indicated concentrations of nDIM (2 µM), nTMZ (5 µM) and dual-loaded (1:2.5 molar ratio) treatment was more effective in inducing apoptosis than treatment with empty vesicles or individually packed nanoformulations, suggesting a synergistic effect of dual-loaded nanoformulation. Further, treatments with individually packed or dual-loaded nanoparticles on C6 cells indicated increased caspase-3 cleavage and Apaf1 intensity, indicating a more pronounced apoptotic effect in dual-loaded nanoformulation. Additionally, PARP cleavage and increased cytochrome C expression demonstrate a discernible rise of apoptosis induction, which is more prominent when dual-loaded nanoparticles was used (**Fig. 4C**). Furthermore, bright field microscopy was used to confirm this differential activity, and our results revealed that treatment with dual-loaded nanoparticles was more effective to decrease both cell numbers and cell viability (**Fig. 4D**).

**Fig. 4.**
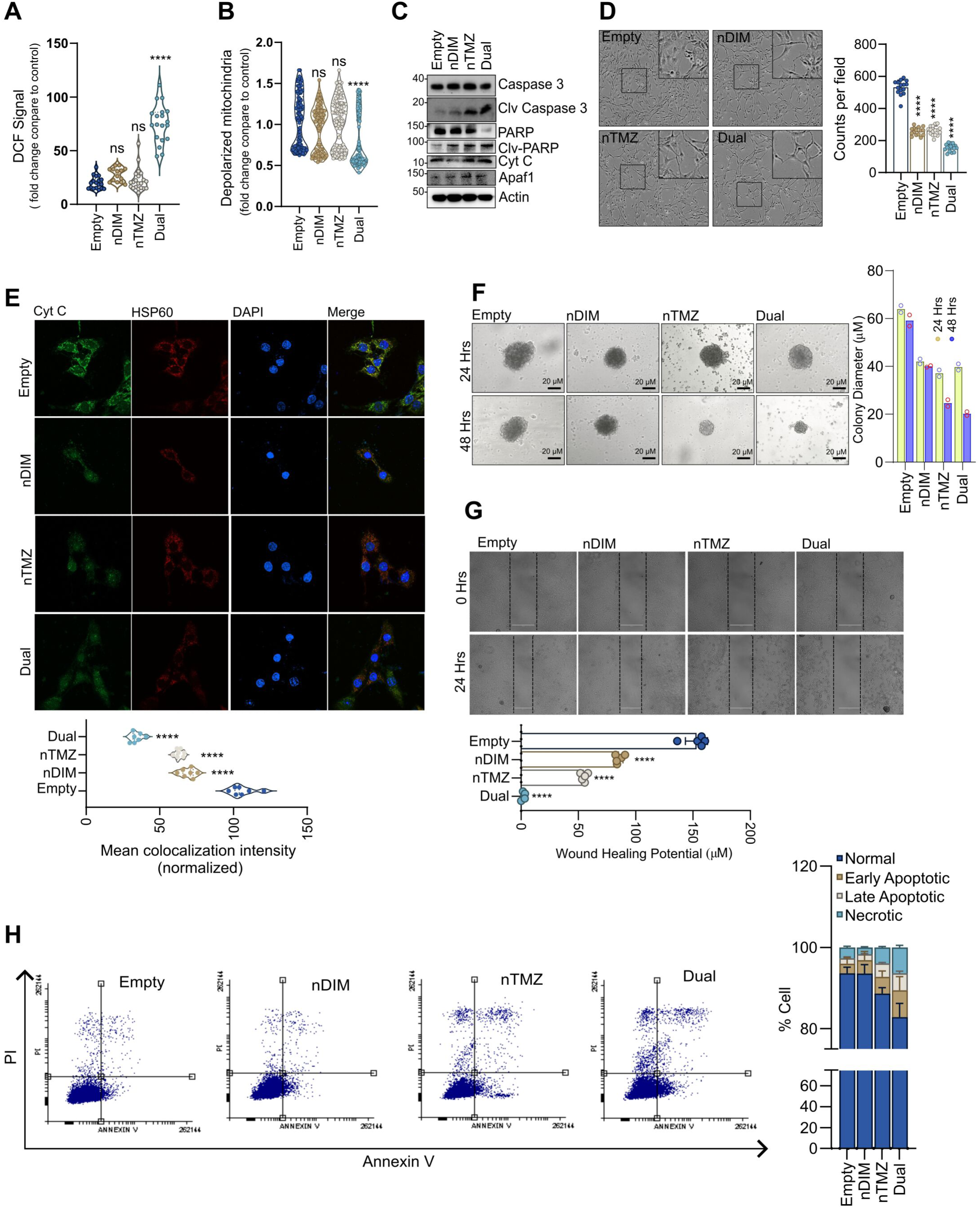
*In vitro* biological evaluation of PLGA packed DIM and TMZ nanoformulations. **A.** Comparative ROS generation. **B.** Flow cytometric analysis for MMP of either empty, nDIM, nTMZ or dual-loaded nanoformulations. **C.** Immunoblot analysis of C6 cells treated with either empty, nDIM, nTMZ or their dual-loaded nanoformulations. **D.** Bright field analysis, **E.** Cytochrome C and HSP 60 colocalization study **F.**, Colony formation assay, **G.** Scratch assay and **H.** percent apoptosis of C6 cells treated with PLGA encapsulated either DIM, TMZ or dual-loaded nanoformulations.

To confirm the initiation of apoptosis, we checked for the release of Cytochrome C (Cyt C) from the mitochondria using confocal microscopy. The diffused fluorescence intensity of Cyt C and HSP60, along with a decreased colocalization signal, confirmed the release of Cyt C from treated C6 cells mitochondria. Additionally, treatment with individual nanoformulations significantly decreased HSP60 and Cyt C colocalization intensity, which was even more prominent when treated with dual-loaded nanoformulation (**Fig. 4E**). To further confirm the apoptotic potential of the nanoformulations, we also conducted scratch and colony formation assays. Treatment of C6 cells with either nDIM, nTMZ or dual-loaded nanoformulations for 24 and 48 hrs resulted in a significant shrinkage in the size of the soft agar colonies, although changes were not prominent after 24 hrs of treatment (**Fig. 4F**). Additionally, the scratch assay revealed a lower wound healing potential, as evidenced by decreased cell migration in the scratched region after 24 hrs of treatment (**Fig. 4G**). Furthermore, Annexin V/PI flow cytometric studies showed that the percentage of apoptotic cell population (including early, late, and necrotic cells) induced by the dual-loaded nanoparticles after 24 hrs of treatment was ∼ 17.5%, compared to ∼6.5% and ∼11.34% for nDIM and nTMZ, respectively. Taken together, all these results suggest that DIM had a better synergistic effect with TMZ in the dual-loaded nanoformulation than in their individually packed nanoformulations (**Fig. 4H**).

### In-vivo efficacy of individually packed and dual-loaded nanoformulations in rat xenografts

To validate the effectiveness of individually packed and dual-loaded nanoformulations *in vivo* xenograft tumor models need to be tested following promising *in vitro* results. Initially, blood serum samples were analyzed for hepatotoxicity and nephrotoxicity in healthy rats who received recommended doses of both native and individually packed DIM and TMZ. Results indicate that rats treated with nanocompounds displayed insignificant changes in ALT, AST, UREA, and creatinine levels compared to their untreated counterparts. Conversely, rats given only unpacked drugs demonstrated significant changes in these parameters, implying toxicity of their native states (**Supplementary Table 1**). CDX tumors were generated in female Sprague Dawley (SD) rats by injecting C6 glioma cells in the flank region. The rats were closely monitored for tumor-related symptoms or complications during the treatment periods. A 30-day treatment with either individually packed or dual-loaded nanoformulations of nDIM and nTMZ near the tumor region significantly reduced tumor growth, while the empty nanoformulation was kept as control. Rats treated with dual-loaded nanoformulation did not display any significant changes related to tumor size, while the control group showed early signs of rapid tumor growth during the treatment period. Furthermore, the weight and volume of tumors were considerably lower in rats treated with dual-loaded nanoformulation than those treated with individually packed molecules and the control groups (**Fig. 5A, B, and C**). The histological appearance of the control tumor tissue sections showed an increased cellular density with prominent nucleoli and a spongy appearance, whereas no such change in tumor morphology was detected in the treated groups. The effectivity of dual-loaded nanoformulation in reducing tumor growth was further analyzed by using immunohistochemical (IHC) analysis. A higher PCNA staining intensity in the control tumor tissue sections indicated an increased uninterrupted proliferation, whereas the reduced expression in the treated groups may be due to the induction of apoptosis. IHC analysis also showed increased activation of cleaved caspase-3 in dual-loaded nanoformulation treated tumors (**Fig. 5D**). The TUNNEL assay further confirmed the induction of apoptosis resulted in more TUNNEL-positive cells in the treated groups. Comparing tissue sections from treated tumor-bearing rats to those from rats treated with empty vesicles, the representative IF images revealed more TUNNEL-positivity in the treated groups. These outcomes demonstrated that tumor cells from treated rats experienced inter-nucleosomal fragmentation of genomic DNA (**Fig. 5E**). The aforementioned results were further supported by IB analysis, where the apoptotic markers like PARP and Caspase 3 were cleaved after being treated with the dual-loaded nanoformulation, as shown by a detectable increase in cleaved caspase-3 and PARP in the treated groups (**Fig. 5F**).

**Fig. 5.**
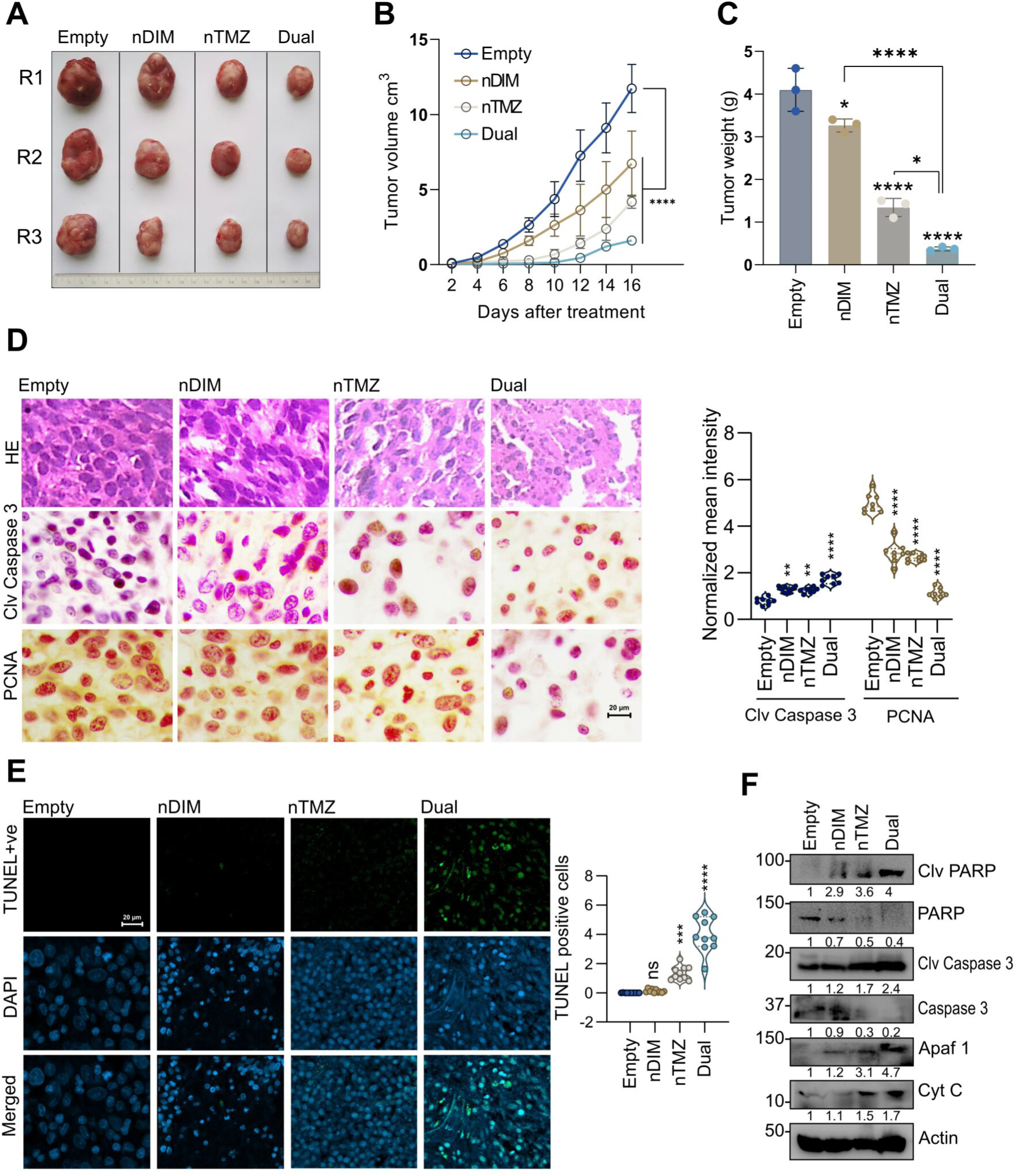
*In vivo* efficacy of different nanoparticles in rat glioma subcutaneous allografts. **A.** Representative images of tumors in the flank region of SD rats after treatment with PLGA encapsulated DIM, TMZ and dual-loaded nanoformulations; empty nanoparticle treatment was kept as control. **B.** Comparison of tumor volumes of different groups. **C.** Tumor weight of different groups, treated and vehicle treated (n=4). Data is represented as Mean±SD. **D.** H&E staining and representative analysis of IHC images of cleaved caspase 3 levels and PCNA expressions, Scattered plot represents mean intensity of PCNA and cleaved caspase 3 of all the groups (Treated and vehicle treated); **E.** TUNNEL assay of different tumor tissues treated with DIM and TMZ nanoparticles (either individual or dual-loaded) showing comparative TUNNEL positive cells. **F.** Immunoblots of either individual packed or dual-loaded nanoformulation treated tumor tissues.

### Efficacy of dual-loaded nanoformulation in rat intracranial orthotopic model in vivo

Using an orthotopic intracranial glioma model in SD rats, we assessed the effectiveness of a dual-loaded nanoformulation. The rats were treated every alternate day with the dual-loaded nanoformulation for 26 days, resulting in significant growth inhibition. In contrast, the control group rats similarly injected with empty nanoparticles displayed tumors that showed no regression. Body weight significantly decreased in the control groups receiving empty nanoparticles, demonstrating that excessive tumor growth interfered with vital physiological functions (**Fig. 6A**). The cellular density and spongy appearance of the tumor tissues were reduced after treatment, according to H&E analysis and manual examination. This data indicates that tumor cells gradually die after receiving dual-loaded nanotherapy. The effectiveness of the dual-loaded nanoformulation in arresting the growth of intracranial tumor cells was further examined using H&E and IHC. We found that the expression of PCNA was decreased and cleaved caspase-3 was increased in the treated groups, which is indicative of the induction of apoptosis, whereas higher PCNA staining intensity and increased cell density were observed in the tumor tissue sections of control groups, indicating a higher uninterrupted proliferation (**Fig. 6B**). Next, TUNNEL assay was used to further confirm this observation. The representative immunofluorescence images revealed more TUNNEL-positive cells in the dual-loaded nanoformulation treated tumor sections compared to the control groups. This outcome demonstrated that tumor cells in treated rats underwent inter-nucleosomal degradation of genomic DNA (**Fig. 6C**). In summary, this study describes the synergistic actions of both DIM and TMZ as PLGA-based dual-loaded nanoformulation after successful delivery to the brain across the BBB resulting in tumor regression (**Fig. 6D**).

**Fig. 6.**
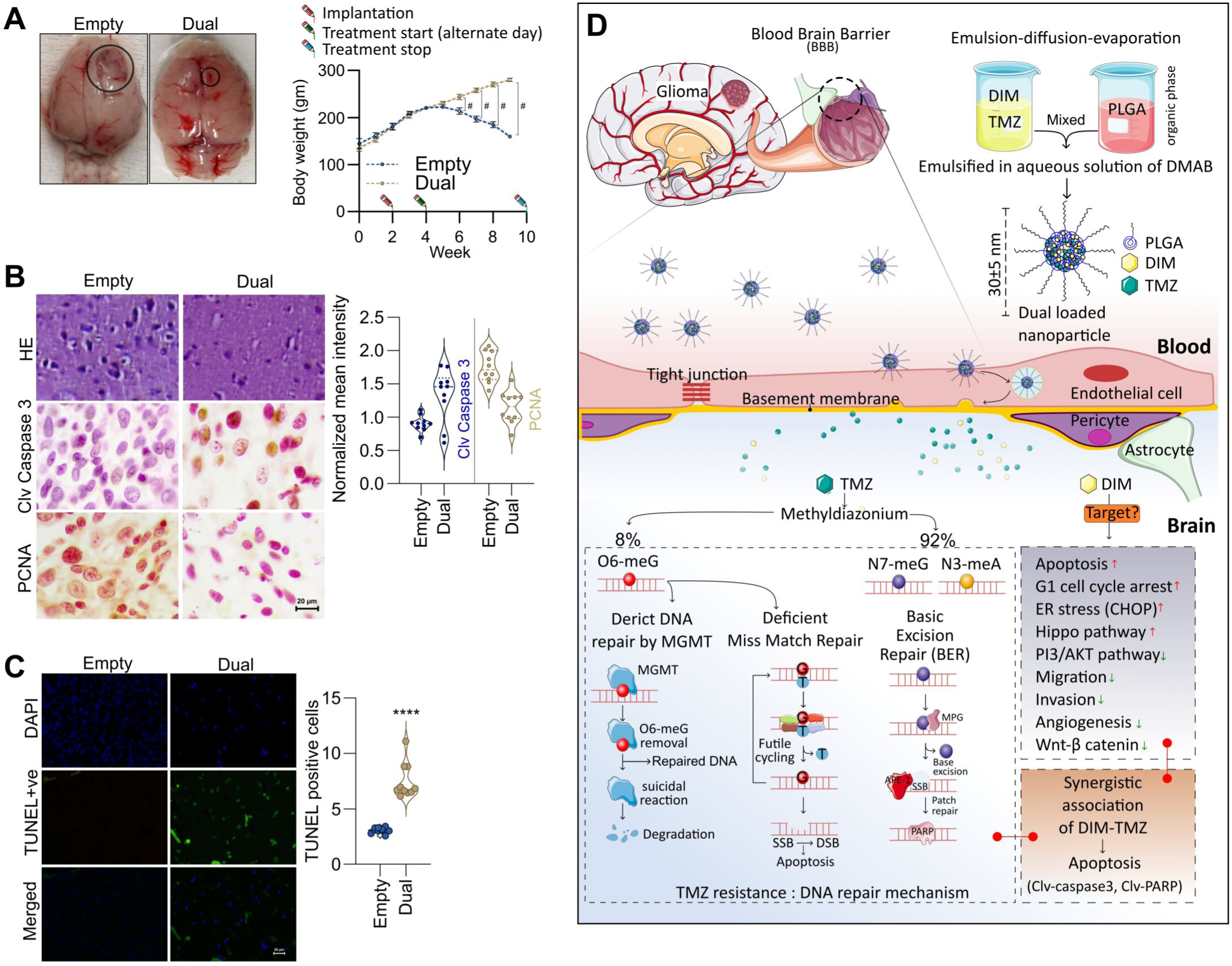
*In vivo* efficacy of dual-loaded nanoparticles in orthotropic/intracranial tumors. **A.** Representative images of dual-loaded nanoformulation or empty vehicle treated rat brain tumors (Orthotopic models). Comparison of body weight of same dual-loaded nanoformulation or empty vehicle treated tumor bearing rats (n=4). Data is represented as mean ±SE. **B.** H&E staining and IHC image analysis of PCNA and cleaved Caspase 3 in vehicle treated and dual-loaded nanoformulation treated tumor tissue sections. Scattered plot represents normalized mean intensities. **C.** TUNNEL-positive cells in empty vehicle and dual-loaded nanoformulation treated groups. **D.** Working model of the dual-loaded nanoformulation and their delivery to brain tumor region across the BBB.

## Discussion

The scientific community continues to challenge the highly aggressive nature of glioma to be the most difficult healthcare issue, and even with standard treatments patient survival is less than 15 months ^38^. The BBB is considered to be the main barrier to glioma treatment, and because of its presence, the majority of chemotherapeutic drugs cannot enter the brain ^39^. Because of this problem, the potential for nanoparticles to penetrate brain parenchyma has emerged as a promising area of glioma research ^40^. The rapid necrosis of tissues and microvascular proliferation produces a semipermeable leaky BBB. Additionally, the EPR effect and damage of the lymphatic systems in the brain parenchyma allow nanoparticles to cross the BBB ^19^.

There are several limitations to the use of chemotherapeutic drugs in treating glioma. As for example, TMZ is commonly used in glioma therapy but after prolonged use, it shows resistance through the involvement of O6-Methylguanine-DNA Methyltransferase (MGMT). But, genotoxic adduct (O6 meG) is neutralized by MGMT and restore DNA integrity by suicidal mechanism in repairing DNA directly ^41^. Generally, DNA repair mechanism involving the DNA mismatch repair (MMR) ^42^ and base excision repair (BER) systems ^43^ contribute to chemoresistance against the drug under treatment. However, DIM, a natural product based molecule, inhibits tumor growth and cause cell cycle arrest in many cancer cells but due to its hydrophobic nature it cannot reach the tumor inside the brain and act on glioma cells. It will also reduce CDK2 activity by upregulating the expression of p21, a molecule accountable for cell cycle arrest. It also inhibits β-catenine and involved in negative regulation of c-Myc, FOS and cycline D1 ^44^. Thus, to improve glioma therapy, in this report we explored the effect of TMZ on C6 glioma cells in combination with DIM (Fig 1A). We observed that both DIM and TMZ have inhibitory effects on glioma cells in a dose dependent manner. The toxic effect of TMZ was more significant on C6 cells than that of DIM when used individually, but based on the Chou-Talalay the strongest synergistic effects of DIM and TMZ were examined and IC50 values were also significantly decreased (Fig 1C). Thus DIM indeed synergistically improve the therapeutic effect of TMZ and reduces TMZ toxicity.

The current study created a novel delivery strategy using brain-penetrating PLGA nanoparticles that encapsulate DIM and TMZ together to improve their solubility, target specificity and efficacy at much lower concentrations to address the current drug delivery challenges for the treatment of glioma. This delivery system can also address the issues of hydrophobicity and low bioavailability as well as enhancing the medications effectiveness. The physicochemical characterization showed that nanoparticle sizes were less than 50 nm and optimized even when loaded with two drugs together. The physical stability of nanoformulations was determined by Zeta potential measurement and found to be easily dispersed in water. There was also no structural change of the loaded compounds in the nanoparticles, as detected by FTIR studies. Additionally, it had been observed that both DIM and TMZ showed sustained release from the nanoparticles, where we used DMAB as a surfactant during preparation to increase the entrapment efficiency and showed significant protective effects against harsh *in vivo* conditions ^26^. DMAB also acts as a stabilizer that leads to dispersed nanoparticles. A stable positive charge of the nanoparticles bound to the glioma cells through electrostatic force that released the drugs slowly in a sustained manner at the specified sites and exhibited perinuclear localization followed by endosomal escape due to the “protein sponge” effect. Apoptotic induction of tumor cells is an important method to observe the effectivity of drug molecules. In this study, it was further observed that small amount of drugs when used in combination compared to single drug, act more effectively on the cells by dissolving and diffusing across the nuclear membrane into the nucleus to ensure organelle damage and be able to induce apoptosis (Fig 4).

Dual-loaded nanoformulation can also efficiently cross the BBB, target brain tumor cells, and achieve increased therapeutic efficiency through their synergistic modes of actions ^45^. When delivered to the tumor site, the dual-loaded nanoparticles showed excellent release profiles and remained stable when observed by Zeta potential measurement (Fig 3). Furthermore, our results on cytotoxic studies showed apoptosis caused *via* caspase-3 dependent pathway (Fig 4C) and produced more TUNNEL-positive cells in treated tumor tissues than untreated. Small amount of DIM and TMZ packed in dual-loaded nanoformulation outperformed compared to the individually packed and unpacked drug molecules (Fig 4). This indicates that the compounds successfully internalized in the PLGA nanoparticles and were able to cross the BBB to enhance the drug delivery inside the tumour microenvironment (TME). Furthermore, apoptosis-mediated cell death was also linked to mitochondrial damage and our results indicated by MMP loss and reactive oxygen species (ROS) generation in treated cells (Fig 4A and 4B). The apoptosis-mediated cell death was more pronounced in the dual-loaded nanoparticles. This combination treatment increases the cytotoxic effect of TMZ and prevents the possible acquired chemoresistance against it. The translational potency of this combinatorial approach, as next-generation glioma therapy, is further demonstrated by the findings showed its more effectiveness than individually loaded nanoformulations.

The post-treatment survival of the tumor-bearing rats was dramatically extended in dual-loaded nanoformulation compared to the treatment-free control groups and treated with individually packed TMZ or DIM. It has been observed that there was a significant inhibition of tumor growth by dual-loaded nanoformulation in both subcutaneous and intracranial tumor-bearing rat models. The increased tumor efficacy through inhibitory effects was due to enhanced blood circulation time and sustained release of the loaded compounds from the nanoparticles for a long time due to the EPR effect ^19, 46^. The counting of TUNNEL-positive cells and IHC studies in tissue sections of untreated *vs* treated tumors of subcutaneous and intracranial models strongly support the successful treatment outcome (Fig 5 & 6). Thus, the novelty of our study points out the treatment as dual-loaded nanoformulation is very promising for further clinical investigations.

In conclusion, nanoformulations of anticancer molecules show promise for glioma therapy due to their improved bioavailability and higher apoptotic effect result in increased survival of intracranial tumor-bearing rats at lower drug doses when compared to their native forms. Therefore, targeted delivery of nanoformulations and combination treatments as dual-loaded nanoformulation would have a better clinical outcome for treating brain cancer patients in the near future.

## Funding

This work is jointly supported by the Department of Science and Technology {NanoMission: DST/NM/NT/2018/105(G); SERB: EMR/2017/000992} and Focused Basic Research (FBR): MLP-142 and HCP-40, CSIR, Govt. of India to Dr. Mrinal K Ghosh. Department of Health Research (DHR: No. 312816 R 12013/10/2017-HR) to Dr. Sibani Sarkar.

## Conflicts of interest

The authors gladly declare no conflicts of interest.

## Acknowledgement

Authors gladly acknowledge Mr. Partha S. Mohanta and Dr. Arijit Bhowmik, Ex-student of Dr. Mrinal K Ghosh for technical help in this study.

## Supplementary materials

**Supplementary Table 1.** Effect of native and nanoformulations treatment in serum ALT, AST, Urea and Creatinine activity in Sprague Dawley rat.

**Supplementary Fig. 1.** Mass spectrometry based confirmation of individually nanoencapsulated DIM and TMZ.

## Notes

### Competing Interest Statement

The authors have declared no competing interest.

