## Supplementary data – Supplementary Figure 1 and Table 1. for "PLGA-based dual-loaded nanoformulation of DIM and TMZ - An advanced clinical strategy for brain cancer treatment in a combinatorial approach"

**A**

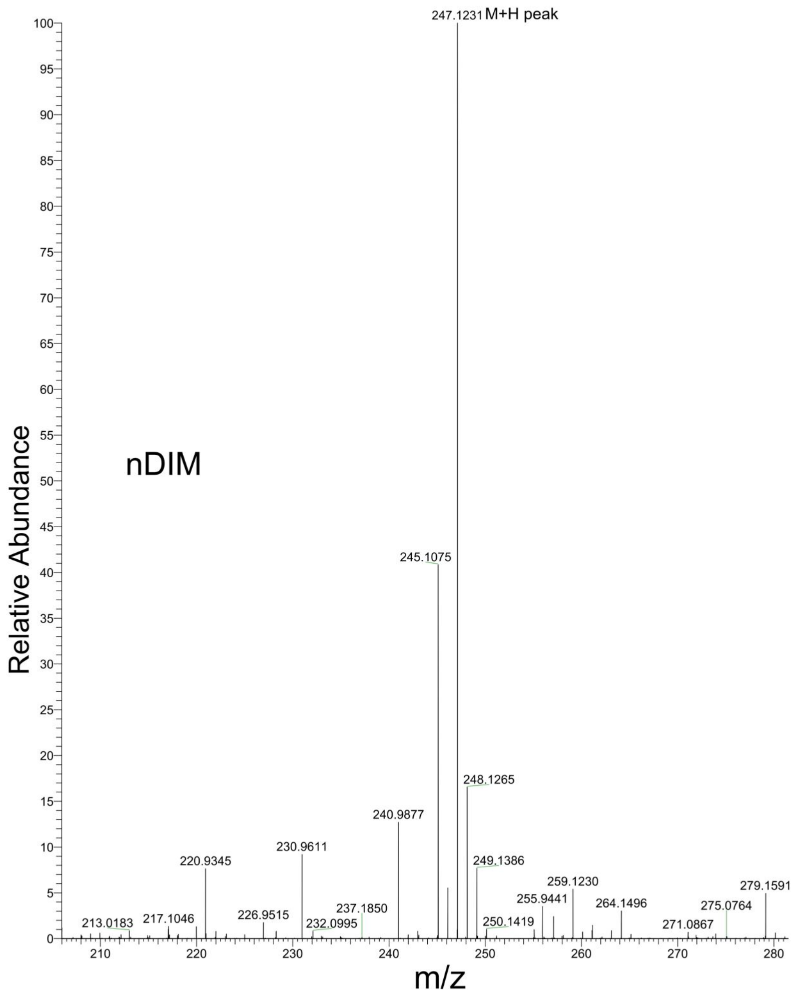


**B**

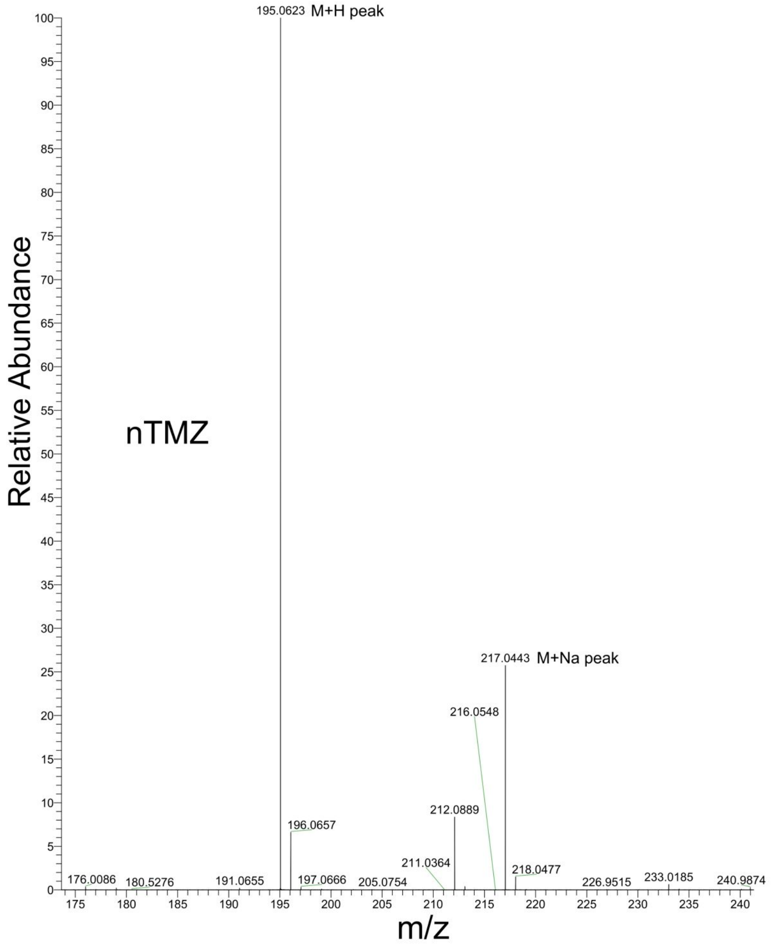

**Supplementary Table 1.** Effect of native and Nano formulations treatment on serum ALT, AST, Urea and Creatinine activity in Sprague Dawley rat.

| Groups | Dose(mg/kg) | ALT (U/ml) | AST (U/ml) | Urea (mg/dl) | Creatinine (mg/dl) |
|---|---|---|---|---|---|
| Vehicle Control | - | 110.70±0.82 | 140.20±0.48 | 7.41±0.043 | 0.93 ±0.042 |
| DIM | 2.5 | 98.25±0.63 | 132.55 ±0.49 | 6.72±0.016 | 0.89 ±0.024 |
| nDIM | 2.5 | 84.13±0.52 | 121.17 ±0.84 | 5.26 ± 0.081 | 0.84 ±0.062 |
| TMZ | 2.5 | 104.77±0.89 | 135.40 ± 0.56 | 6.91±0.042 | 0.88 ±0.058 |
| nTMZ | 2.5 | 76.19 ± 0.63 | 112.26 ±0.47 | 4.91±0.077 | 0.82±0.049 |
| Dual | 2.5 | 68.26 ± 0.47 | 101.24±0.63 | 3.86±0.035 | 0.75 ±0.036 |

Data were represented as mean ± SD, n = 6. *P<0.05 significantly different from vehicle control group.
